# Upregulation of Somatostatin Receptor Type 2 in a Receptor-Deficient *In Vivo* Pancreatic Neuroendocrine Tumor Model Improves Tumor Response to Targeted ^177^Lu-DOTATATE

**DOI:** 10.1101/2022.04.25.489401

**Authors:** Rupali Sharma, Bhargav Earla, Kwamena Baidoo, Martha A. Zeiger, James P. Madigan, Freddy E. Escorcia, Samira M. Sadowski

## Abstract

**Purpose:** The goal of this study was to test whether histone deacetylase inhibitors (HDACis) restore somatostatin receptor type 2 (SSTR2) expression in models of high-grade pancreatic neuroendocrine tumors (PNETs), thereby facilitating effective treatment with ^177^Lu-DOTATATE therapy.

**Methods:** To assess tumor grade correlation with SSTR2 expression, we assessed human SSTR2 promoter methylation and expression levels in 96 NIH patient samples and merged the GSE149395 and GSE117852 datasets. We used three NET cell lines (QGP-1, BON-1, GOT-1) characterized by variable SSTR2 expression profiles for functional *in vitro* studies using HDACis. Finally, the QGP-1 xenograft mouse model, with low basal SSTR2 expression, was used to analyze the therapeutic efficacy of combined HDACi and ^177^Lu-DOTATATE therapies.

**Results:** Human PNET SSTR2 promoter methylation showed a significant positive correlation with higher tumor grades (*P* = 0.000014). We also found a significant negative correlation (*P* < 0.0001) between SSTR2 promoter methylation and SSTR2 expression in three NET cell lines. *In vitro*, SSTR2 expression increased significantly in BON-1 and QGP-1 cells at 48 and 72 hours in a dose-dependent fashion using two different HDACis, valproic acid and CI-994. *In vivo* studies demonstrated a significant increase in ^177^Lu-DOTATATE tumor uptake in QGP-1–engrafted mice after 10 days of CI-994 pretreatment (*P* = 0.0175). Treatment with ^177^Lu-DOTATATE reduced tumor size in mice pretreated with CI-994 compared to ^177^Lu-DOTATATE alone (at 15 days, *P* = 0.0028).

**Conclusion:** HDACis increase SSTR2 surface expression in models of high-grade, SSTR2-deficient PNETs. This approach has the potential to improve tumor response to targeted therapy with ^177^ Lu-DOTATATE in patients with receptor-negative, metastatic PNETs.

**Translational Relevance Statement:** Pancreatic neuroendocrine tumors (PNETs) express high levels of somatostatin receptor type 2 (SSTR2), a unique target for both tumor imaging and therapy. Unfortunately, high-grade PNETs lose SSTR2 surface expression and thus become ineligible for SSTR2-targeted ^177^Lu-DOTATATE peptide receptor radionuclide therapy (PRRT). Restoring SSTR2 expression through the reversal of inhibitory epigenetic gene silencing mechanisms has the potential for improving tumor responsiveness to PRRT. We demonstrate that histone deacetylase inhibitors (HDACis) upregulate SSTR2 surface expression in three NET cell lines *in vitro*. In an *in vivo* PNET xenograft model with low basal SSTR2 expression, our studies validate a significantly higher tumor uptake of SSTR2-targeted ^177^Lu-DOTATATE in animals pretreated with HDACis compared to controls. Furthermore, we show that this higher tumor uptake results in significant anti-tumor response when compared to standard PRRT alone. Our preclinical results thus provide a rationale for utilizing HDACi pretreatment to improve targeted radionuclide therapy in patients with SSTR2-negative, metastatic PNETs.

## Introduction

While gastro-entero-pancreatic neuroendocrine tumors (GEP-NETs) are rare, their incidences in the United States have increased 6.4-fold over the last four decades (1). Overall survival in high-grade, metastatic NETs is less than 25% (2–4). High-grade, poorly differentiated tumors are associated with poor prognoses (5) and are characterized by a loss of somatostatin receptor type 2 (SSTR2) expression (6). These SSTR2-negative tumors are refractory to targeted therapies, respond poorly to chemotherapy (streptozosin- or platinum-based) (7), and have no effective treatments available. Our prior work resulted in the approval of the positron emission tomography (PET)-imaging agent, ^68^Ga-DOTATATE (6, 8). This agent is a high-affinity ligand that binds SSTR2, which has not only revolutionized the diagnosis of low-grade NETs, but also allows for peptide receptor radionuclide therapy (PRRT) with ^177^Lu-DOTATATE (9). However, because high-grade NETs harbor negative or low levels of SSTR2, PRRT is not effective in these patients. Thus, herein we propose to render these receptor-negative NETs treatable by pharmacologically increasing SSTR2 expression.

Epigenetic regulation is one of several mechanisms used by cells to modulate gene expression. DNA methylation and specific histone modifications can lead to transcriptional inactivation of tumor suppressor genes and are directly correlated with tumorigenesis of GEP-NETs (10). To date, no genomic alterations have been found in the *SSTR2* gene of NETs. Additionally, NETs have a low mutation burden, and increased DNA methylation correlates with poor patient outcomes. Thus epigenetic regulation may play a dominant role in the oncogenesis and progression of NETs, and account for the lack of SSTR2 in high-grade NETs (11). A few preclinical studies have shown that epidrug treatment significantly increased SSTR2 expression in cell lines with low basal expression (12–14). However, how this translates to *in vivo* animal models has not been previously demonstrated.

In the current study, we first confirm increase in SSTR2 expression in three GEP-NET cell lines (QGP-1, BON-1, and GOT1) using targeted epidrug treatment (the histone deacetylase inhibitors [HDACis] valproic acid [VPA] and tacedinaline [CI-994] and the DNA methyltransferase inhibitors [DNMTis] azacitidine and decitabine). To validate the translational potential of these findings, we demonstrate that these epidrugs can increase *in vivo* uptake of ^177^Lu-DOTATATE and for the first time confirm the therapeutic efficacy of PRRT with ^177^Lu-DOTATATE in a preclinical model of SSTR2-deficient pancreatic NETs (PNETs).

## Materials and Methods

### Compounds

HDACis, VPA and CI-994, and DNMTi, decitabine, were purchased from Sigma-Aldrich and Azacitidine from Selleckchem. CI-994 and decitabine were dissolved in DMSO (Sigma-Aldrich), VPA dissolved in water and, all stored at –20°C. Concentrations of each inhibitor were calculated based on previously published data, with concentrations below the maximum tolerated dose (MTD) used in human trials (15–19). ^177^LuCl_3_ was obtained from MURR (MU Research Reactor, University of Missouri, Research Park Drive Columbia, MO). ^177^Lu-DOTATATE was synthesized as described (20).

### EPIC/850k methylation array and RNA sequencing in human samples and cell lines

GEP-NETs samples, collected from patients under a prospective protocol at NIH (IRB approved, 09-C-0242), were scored using the World Health Organization (WHO) classification system (21). 850K/EPIC array data in a cohort of 96 samples of NETs from NIH were analyzed using R-based platforms (22). The raw β values were extracted using the mini package and underwent functional normalization. Probes located in promoter regions of SSTR1-5 were interogated. Methylation levels among different tumor grades were analyzed using the two-tailed *t* test. The Chan *et al.* dataset (23), GSE117852, containing 32 human PNET samples, was analysed using Spearman’s method, with the *P*-value based on the rank of the expression. 850K/EPIC array analysis and RNA-sequencing were performed in QGP-1, BON-1, and GOT-1 cells after a 4-day treatment with VPA and azacytidine. Normalized methylation data were evaluated in R using Champ package functions (champ.norm) and the beta-mixture quantile normalization method.

### Cell culture

Three NET cell lines were used: QGP-1, BON-1, and GOT-1. BON-1 was established from a lymph node metastasis from a PNET patient, provided by Mark Hellmich (University of Texas Medical Branch at Galveston) (24). The cell lines were maintained in a 1:1 mixture of high-glucose DMEM and Ham’s F-12 nutrient mix supplemented with 10% FBS and 1% penicillin-streptomycin. QGP-1 was established from a somatostatin-producing pancreatic islet cell carcinoma (25), purchased from the Japanese Collection of Research Bioresources cell bank and maintained in glutamine-containing RPMI-1640 medium with 10% FBS and 1% penicillin-streptomycin. The GOT-1 cell line was established from a liver metastasis of a primary intestinal NET and provided by Ola Nilsson (Sahlgrenska Cancer Center, University of Gothenburg, Sweden) (26, 27). The cells were maintained in glutamine-containing RPMI-1640 medium with 10% FBS, 5 µg/mL insulin, 5 µg/mL transferrin, 100 IU/mL penicillin, and 100 μg/mL streptomycin. Cells were maintained at 37°C in a humidified environment at 5% CO_2_. Cell lines were authenticated by STR analysis via Labcorp’s human cell line authentication service, and were routinely tested for mycoplasma infection using the MycoAlert PLUS Mycoplasma Detection Kit (LT07-705, Lonza).

### RNA extraction and real-time reverse transcription PCR

For mRNA analysis, cells were plated in 6-well plates and total RNA was isolated from the cells using RNeasy Plus Mini kit (#74106, Qiagen). The RNA concentrations were determined using a NanoDrop 1000 spectrophotometer (Thermo Fisher Scientific) and 600 ng of RNA were used to synthesize cDNA using a high-capacity cDNA reverse transcription kit (Thermo Fisher Scientific). The sequences of the PCR primers used in this experiment are as follows: SSTR2 Hs00265624_s1 and β-actin Hs01060665_g1. Real-time quantitative PCR was performed in triplicate on a Thermo Fisher QuantStudio 5 Real-Time PCR System (Thermo Fisher Scientific). Target gene expression was normalized to β-actin, and the ΔΔCt method was used to calculate relative gene expression.

### Cell surface expression of SSTR2

To analyze cell membrane expression of SSTR2, cells were harvested, washed, and counted. QGP-1, BON-1, and GOT-1 cell lines were treated for up to 5 days with HDACi. After treatment, the cells were fixed in 4% PFA and resuspended in flow buffer (0.5% BSA in sterile PBS). Then, 1×10^6^ cells per replicate were stained in a dark room for 30 minutes at room temperature with an anti-SSTR2 (IC4224G, 1 µg/5µL) antibody (R&D Systems). Cells were then analyzed by flow cytometry using a BD LSR-Fortessa analyzer (BD Biosciences) for SSTR2 surface expression. Data were analyzed using FlowJo V10.6.2 (Tree Star, Inc.).

### Western blot analysis

Cells were treated, lysed with GPCR Extraction and Stabilization Reagent buffer (#A43436, Thermo Fisher Scientific). Cell lysates were rotated at 4°C for 20 minutes and then briefly sonicated twice. Protein concentrations were determined using a Micro BCA Protein Assay Kit (#23235, Thermo Fisher Scientific). Fifty µg of protein lysate were resolved on Bolt^TM^ 4–12% Bis-Tris Plus Gels (#NW04120BOX, Life Technologies) and electrophoresed at 200V for 30 minutes. Gels were transferred to PVDF membranes using the iBlot2 dry transfer method (# IB21001, Life Technologies). Membranes were blocked in 3% BSA/5% milk in 1X Tris buffer saline/0.05% Tween 20 (1X TBST) for 1 hour at room temperature followed by primary antibody incubation overnight, at 4°C, washed three times for 5 minutes in 1X TBST, then incubated with a secondary antibody for 1 hour at room temperature. Membranes were then incubated with a chemiluminescent substrate (#34577, SuperSignal West Pico Chemiluminescent Substrate, Thermo Fisher Scientific) and imaged on the BIO-RAD Chemidoc^TM^ Imaging System (Hercules). Densitometry was performed with NIH ImageJ software (2.0.0) (28), with all protein signal intensities normalized to GAPDH or β-actin.

### SSTR2-specific antibody validation through SSTR2 stable knockdown

(Supplementary Fig. 1A and B): To stably knock down expression of human SSTR2, two separate pLKO.1-puro lentivirus plasmid-based shRNAs targeting the sequences TGAAGACCATCACCAACATTT (#64) and CCCTTCTACATATTCAACGTT (#87) (human MISSION shRNA clone ID: TRCN0000358164, SSTR2 #64-shRNA, and human MISSION shRNA clone ID: TRCN0000014387, SSTR2 #87-shRNA, Sigma-Aldrich) were employed. Additionally, a non-targeting shRNA plasmid (NT-shRNA) that targets no known human sequence was used as a control. A primer containing the target sequence (CTGGTTACGAAGCGAATCCTT), along with a stem-loop followed by the reverse target sequence, was annealed to a complementary primer and inserted into the EcoRI and AgeI sites of pLKO.1-puro (#8453, Addgene). Lentiviral particles were produced *via* Lipofectamine 2000 (Invitrogen)-mediated triple transfection of 293T cells with the respective pLKO.1-puro shRNA plasmids, along with the lentiviral envelope plasmid (pMD2.G, #12259, Addgene) and the lentiviral packaging plasmid (psPAX2, #12260 Addgene). BON-1 target cells were transduced with shRNA-containing lentiviral particles in the presence of 8 μg/mL Polybrene (Invitrogen), and stable cells were selected using 2 μg/mL puromycin (Invitrogen). Efficiency of SSTR2 mRNA knockdown was determined using qRT-PCR. Western blot analysis (described above) of control and SSTR2-specific, stable knockdown BON-1 cell lysates was used to validate the efficacy and utility of a commercially available, SSTR2-specific antibody, M01689 (Boster Bio).

### Mouse model

The animal protocol for this study was approved by the NCI/CCR Animal Care and Use Committee. QGP-1 or BON-1 cells (5×10^6^ cells) were injected subcutaneously into both flanks of five- to seven-week old athymic nude mice (Jackson Laboratory). The resulting tumors were further subcutaneously implanted into five-to-seven-week-old athymic nude mice (Jackson Laboratory). This was done to create tumors similar in composition and size (∼1-mm tumors were subcutaneously implanted in each flank, resulting in homogeneous tumors after 3–4 weeks). Only tumors from generations 1 to 3 were utilized. Tumor size was monitored at the beginning and end of the treatments using a Vernier caliper. At initiation of treatment, the tumors ranged from 5 to 7 mm in size. The mice were euthanized if the tumors exceeded 2 cm in diameter, exhibited impeded movement, or if there were signs of breathing difficulty at any point in the study. The mice were randomly grouped into control and treatment groups and then treated *via* intraperitoneal (i.p.) daily injections with either VPA (300 mg/kg; P4543; Sigma Aldrich) dissolved in water, CI-994 (5mg, 7.5mg and 10mg/kg; C0621; Sigma Aldich) dissolved in 0.1% DMSO, or 0.1% DMSO as a control group.

### Cell surface expression of SSTR2

#### Flow cytometry

For flow cytometry analysis, mice with xenografts of BON-1 or QGP-1 tumors were treated with DMSO, VPA (300 mg/kg), or CI-994 (10 mg/kg). At the end of the treatment period, blood was collected by cardiac puncture and the tumors were harvested for FACS analysis. The tumors were dissociated using a human tumor dissociation kit (#130-095-929, Miltenyi Biotech). The cells were then counted and 1×10^6^ cells stained with an Alexa Fluor 488-conjugated anti-SSTR2 (1µg/5µL) antibody (IC4224G, R&D Systems) in a dark room for 30 minutes at room temperature. These cells were analyzed for surface expression of SSTR2 by flow cytometry using a BD LSR-Fortessa analyzer (BD Biosciences). An appropriate Alexa Fluor 488-conjugated mouse IgG2A antibody (1µg/5µL) (IC003G, R&D Systems) was used as an isotype control. Data were analyzed using FlowJo V10.6.2 (Tree Star, Inc.).

### Immunofluorescence

Immunofluorescence (IF) staining was performed on commercially available pancreas tissue slides (Islet Cell Tumor NBP2-77922, Novus Biologicals), as a positive control for SSTR2, and on formalin-fixed, paraffin-embedded mouse tumor samples. Slides were deparaffinized and hydrated using graded alcohols and distilled water, followed by antigen retrieval at pH 6.0 at 90℃ for 20 minutes. Slides were then blocked with a blocking serum. Next, the slides were incubated with the primary antibody for SSTR2 (UMB1)(ab134152;1:25; Abcam) and a secondary antibody, Alexa Fluor 594 (A11037;1:500, Invitrogen). For Pan-acetylated H3 primary antibody (06-599;1:500; EMD Millipore) and secondary antibody (AF594; 1:100, Thermo Fisher Scientific). For γH2AX primary antibody (05-636-1;1:800; EMD Millipore) and secondary antibody (AF488;1:100, Thermo Fisher Scientific). Finally, the slides were mounted using ProLong Gold Antifade Mountant with DAPI (P36935, Invitrogen). All incubations were carried out at room temperature using 1X TBST as washing buffer. Zeiss LSM −8800 Confocal microscope was used for images.

### Biodistribution studies

All living mice were euthanized 24 hours post-^177^Lutetium (Lu)-DOTATATE injection (2 MBq/mouse). In addition to the tumors, the following organs were harvested: heart, lungs, liver, spleen, kidneys, stomach, small and large intestine, muscle, and femur. The counts per minute (CPM) readings, obtained using a 2480 automatic gamma counter (Perkin Elmer), were normalized by tumor weight and biodistribution data were presented as the percentage of the injected dose per gram of tissue (%ID/g).

### Peptide receptor radionuclide therapy

To evaluate the therapeutic efficacy of the treatments, the radionuclide ^177^Lu was labelled with DOTATATE (#H-6318005; Bachem). Both BON-1 and QGP-1 tumor mouse models were pretreated with either CI-994 or DMSO for 10 days. The mice were then randomized into the following treatment groups: 1) saline; 2) ^177^Lu-DOTATATE (30 MBq/mouse); 3) CI-994 followed by saline; and 4) CI-994 followed by ^177^Lu-DOTATATE (30 MBq/mouse). Five minutes prior to ^177^Lu-DOTATATE administration, D-lysine hydrochloride (35 mg/mouse) (#243080010, Thermo Fisher Scientific) was given i.p to block kidney uptake of and prevent nephrotoxicity from ^177^Lutetium (29). The tumor burden in each mouse was monitored twice a week. The tumor burden was calculated using the formula tumor volume = (length×width^2^)/2.

### Statistical analysis

GraphPad Prism 8.1 (GraphPad Software, Inc.) software was used for data analysis. Two-way ANOVA was used to analyze qPCR and flow cytometry data to determine SSTR2 expression differences between the treatment groups. A two-tailed, unpaired Student’s t-test was used for comparisons between two groups. One-way ANOVA with a *post hoc* Tukey’s test was used for comparisons between more than two groups. Two-way ANOVA with interaction effects was performed at different timepoints for the therapeutic study. Data are presented as mean ± SEM. A *P*-value of < 0.05 was considered significant. Error bars show SEM.

## Results

### SSTR2 promoter methylation varies according to tumor grade in GEP-NET patients

We determined the methylation levels of 12 CpG methylation sites at the SSTR2 promoter region in 96 NIH patient samples and correlated those levels with the tumor grades. Higher tumor grades presented with increased methylation levels, as shown in one of the CpG methylation sites (cg19129425) (*P* < 0.05) (Fig. 1A), indicating closed chromatin with suppressed gene expression in high-grade tumors. The International Cancer Genome Consortium (ICGC) (450K EPIC array) and NIH data (850K EPIC array) sets were merged to examine tumor grades and methylation levels. There was a visible difference in the merged data as both have only four overlapping CpG methylation sites for SSTR2. Significant differences were observed in the methylation levels in grade 1 and 2 tumors (*P* = 0.034), but not with grade 3 tumors. Perhaps this is due to the small sample size for grade 3 tumors (very rare), which makes it difficult to obtain statistically relevant information. Analysis of the GSE 117852 dataset of 32 PNET samples showed a significant negative correlation between *SSTR2* gene expression and mean SSTR2 promoter methylation levels (Fig. 1B), indicating low mean promoter methylation levels (open chromatin) in tumor samples exhibiting elevated SSTR2 expression.

**Figure 1.**
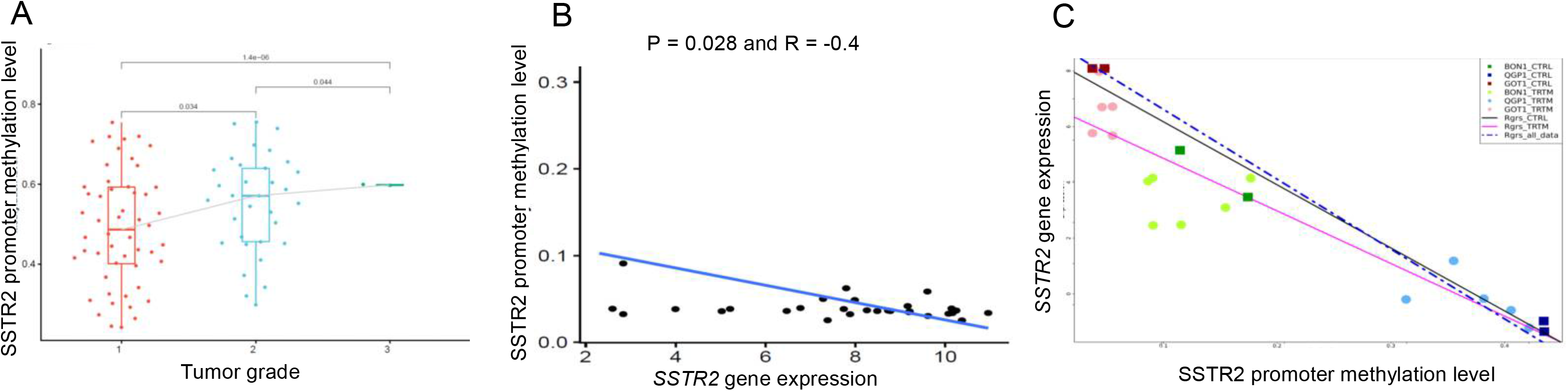
SSTR2 promoter methylation levels in GEP-NET patients and three NET cell lines. **A,** An NIH cohort of 96 patients with GEP-NETs was analyzed based on tumor grade and methylation level. Those patients with higher-grade tumors showed increased methylation levels at the CpG methylation sites of the SSTR2 promoter. Significant changes were observed in the methylation levels between grades 1 and 2 (*P* = 0.034), grades 2 and 3 (*P* = 0.044), and grades 1 and 3 (*P* = 0.000014). **B,** The GSE117852 dataset of 32 patients with PNETs was analyzed for gene expression and mean SSTR2 promoter methylation levels, using the Spearman method to calculate the *R* and *P*-value. **C,** Methylation levels and SSTR2 expression were measured before and after treatment with VPA and azacytidine (TRTM) in three NET cell lines (QGP-1, BON-1, GOT-1). The controls are untreated for comparison (CTRL). The X-axis represents the degree of methylation (0–1), adjusted graphically for optimal visual display. The most significant changes were observed at CpG island cg14232289 (*P* < 0.0001).

### SSTR2 promoter methylation correlates with SSTR2 expression and segregates according to GEP-NET cell line type

We determined the methylation levels of 27 CpG methylation sites at the SSTR2 promoter regions and correlated those levels with SSTR2 expression in three NET cell lines. The NET cell lines demonstrated variable SSTR2 expression, and a significant negative correlation between the level of SSTR2 expression and DNA methylation was found in 11 CpG sites (cg14232289: *P* < 0.0001, Fig. 1C). Significant differences were observed across all three cell lines in their baseline SSTR2 methylation/expression levels and segregated according to cell line types (Fig. 1C, baseline expression = rectangular dots). VPA- and azacytidine-induced methylation changes were seen in the QGP-1 cells, with concomitant changes in SSTR2 expression (Fig. 1C, blue circles). Overall, QGP-1 cells showed the lowest baseline expression of SSTR2 when compared to BON-1 and GOT-1 cells, with correspondingly higher SSTR2 promoter CpG methylation.

### HDACis VPA and CI-994 upregulate SSTR2 expression

Our inhibitor studies (Fig. 1C) demonstrated that treatment with an HDACi assisted in decreasing SSTR2 promoter CpG methylation levels. An important feature of epigenetic regulation is the interconnectedness of disparate epigenetic features and their enzymatic proteins (30) (i.e., separate epigenetic marks laid down by one epigenetic enzyme may interfere in the localization of a different enzyme for an entirely different epigenetic mark (31)). With this in mind, we chose to continue our epigenetic-based studies using HDACis for the potential double-positive effect on gene transcription by both directly interfering with the removal of histone acetylation post-translational marks and interfering with DNA CpG methylation (a negative regulator of transcription) (32).

We found upregulation of *SSTR2* gene and protein expression levels after treatment with the epidrugs VPA and CI-994 in all three cell lines in a dose- and time-dependent manner. Protein expression studies were performed using an SSTR2 antibody that was validated by SSTR2 knockdown experiments (Supplementary Fig. 1A and B). In BON-1 cells, both VPA and CI-994 treatment at 48 and 72 hours yielded a significant, dose-dependent increase in SSTR2 expression (Fig. 2A and B). In QGP-1 cells, VPA treatment yielded a significant increase in SSTR2 expression at 72 hours (Fig. 2C) and CI-994 treatment at 48 hours compared to the controls (Fig. 2D). In GOT-1 cells, lower treatment concentrations of CI-994 significantly increased SSTR2 expression at both 48 and at 72 hours (Fig. 2E). We found a significant 1.4- and 2.8-fold increase in total SSTR2 protein expression with VPA and CI-994, respectively, with a concomitant increase in histone acetylation in the BON-1 cell line at 48 hours, confirming the target and mode of action of the drugs (Fig. 2F). Similarly, in the QGP-1 cell line, a 2.2- and 3.5-fold increase was seen with both HDACis (Fig. 2G). Graphical representation of the fold changes in the BON-1 and QGP-1 cell lines are shown in Fig. 2H.

**Figure 2.**
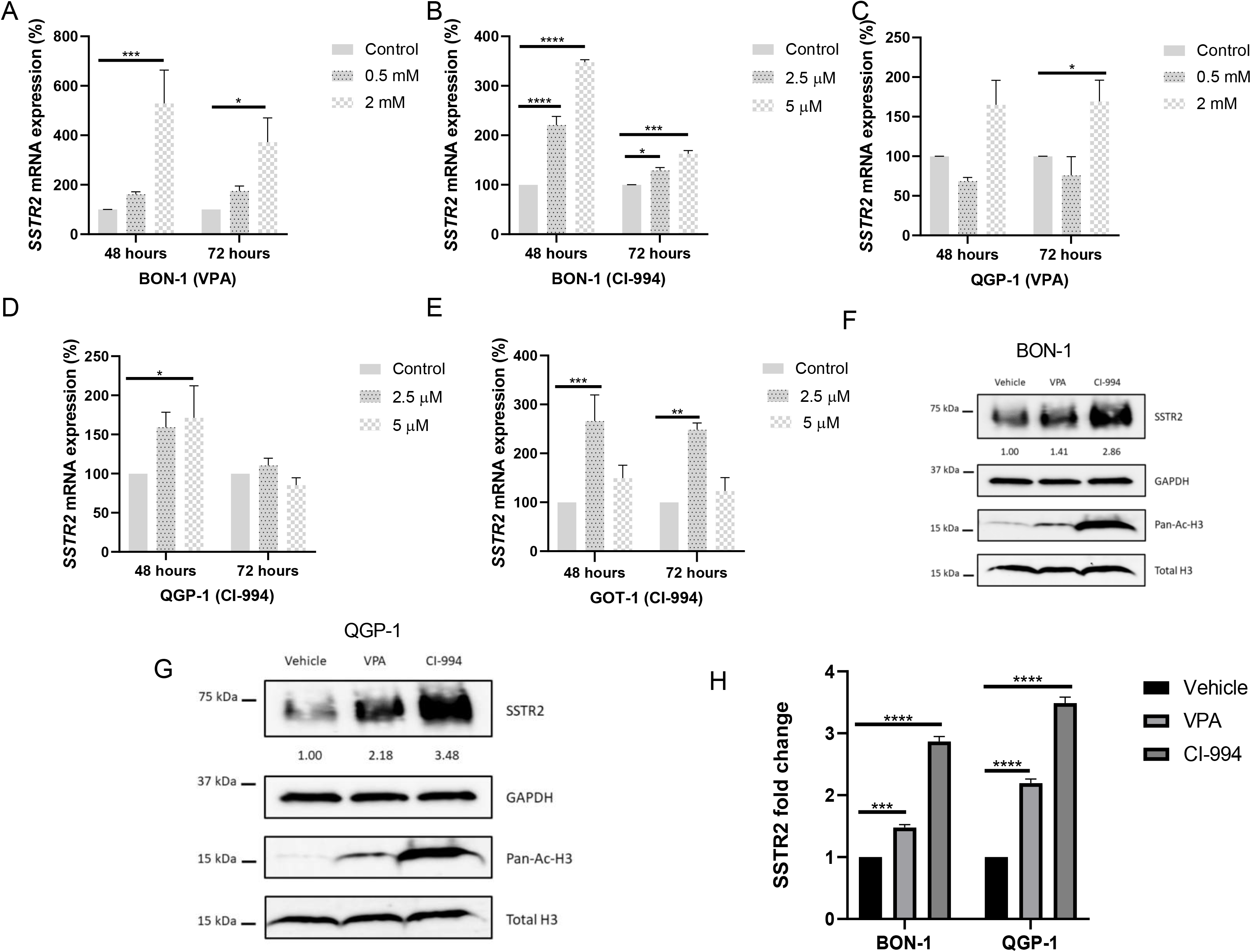
HDACis increase SSTR2 transcription and protein expression. **A,** BON-1 cells treated with VPA at 2 mM for 48 hours (*P* = 0.0001) and 72 hours (*P* = 0.0144) significantly increased SSTR2 mRNA expression. **B,** BON-1 cells treated with CI-994 at 2.5 µM for 48 hours (*P* =0.0001) and 72 hours (*P* = 0.03), and at 5 µM for 48 hours (*P* = 0.0001) and 72 hours (<u>P</u> = 0.0001) had a significant increase in SSTR2 expression. **C,** QGP-1 cells treated with VPA at 2 mM for 72 hours had a significant increase in SSTR2 expression (*P* = 0.0379). **D,** QGP-1 cells treated with CI-994 at 5µM for 48 hours (*P* = 0.0392) had a significant increase in SSTR2 expression. **E,** GOT-1 cells treated with CI-994 at 2.5 µM for 48 hours (*P* = 0.0008) and 72 hours (*P* = 0.0022) showed a significant increase in SSTR2 expression. **F,** HDACis VPA and CI-994 increased SSTR2 protein expression in BON-1 cells 1.4- and 2.8-fold, respectively, with a concomitant increase in H3-histone acetylation at 48 hours. **G,** HDACis VPA and CI-994 increased SSTR2 protein expression in QGP-1 cells 2.18- and 3.48-fold, respectively, with a concomitant increase in histone acetylation at 48 hours. **H,** Graphical representation of the fold change in total SSTR2 protein expression in BON-1 and QGP-1 cells. Data are expressed as mean ± SEM, (*n* = 3). \**P* < 0.05, \*\**P* < 0.01, \*\*\**P* < 0.001, \*\*\*\**P* < 0.0001.

### HDACi treatment increases SSTR2 surface expression in three NET cell lines

At 72 hours, using flow cytometry, we found a significant dose- and time-dependent increase in SSTR2 surface expression in BON-1 cells with both VPA and CI-994 treatment (Fig. 3A and B), and in QGP-1 cells treated with VPA (Fig. 3C). The increase in cell surface expression in QGP-1 with CI-994 was not significant (Fig. 3D); however, when investigating the long-term effects of a 5-day treatment, we found a significant increase in SSTR2 surface expression in CI-994–treated QGP-1 cells (Fig. 3E). At 72 hours, we found a significant increase in SSTR2 surface expression in CI-994–treated GOT-1 cells (Fig. 3F). These results show that HDACi treatment increases plasma membrane SSTR2 in NET cell lines.

**Figure 3.**
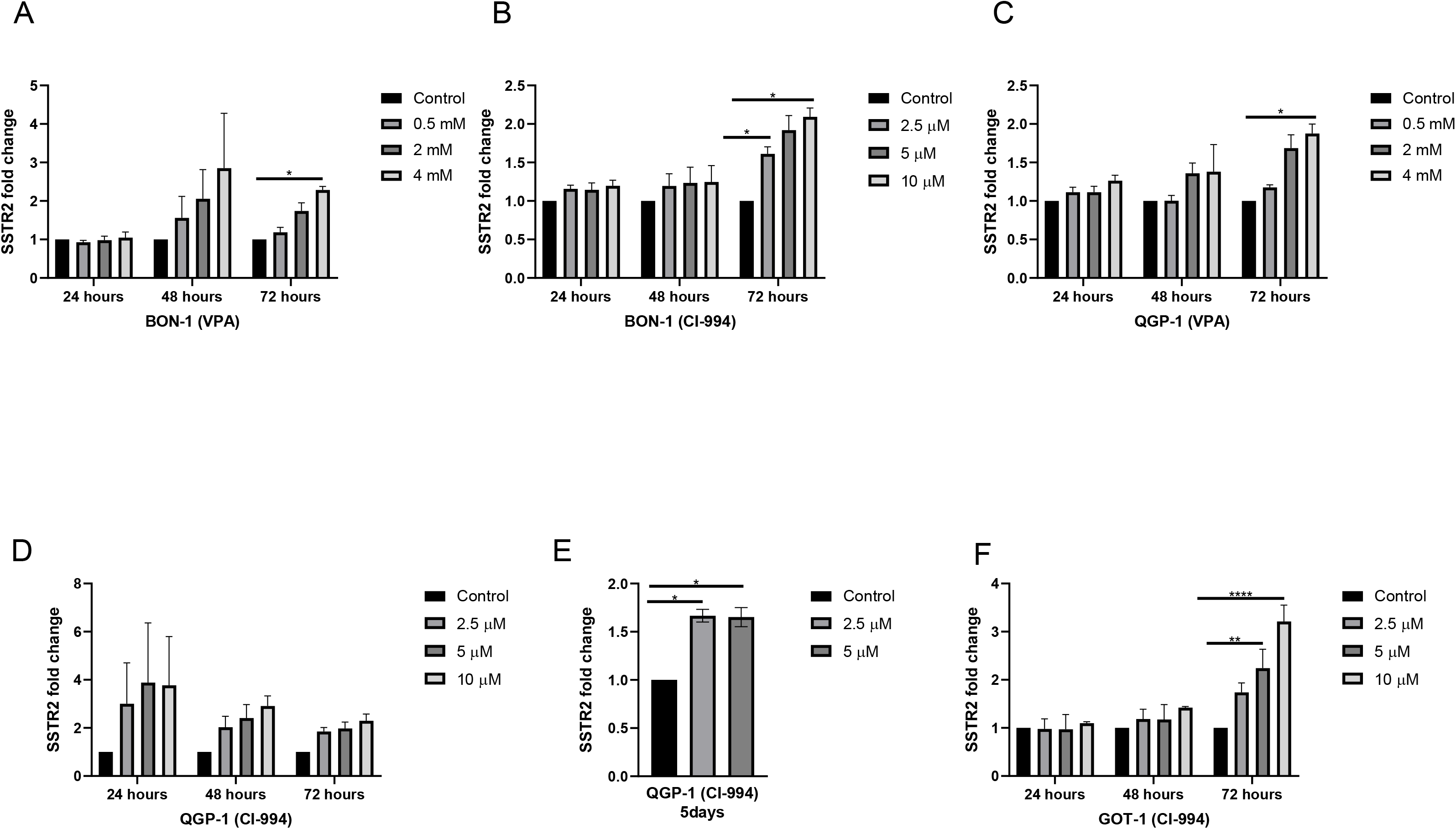
HDACis upregulate SSTR2 surface expression in three NET cell lines. **A** and **B,** SSTR2 surface expression in BON-1 cells was determined by flow cytometry. Cells treated with both VPA at 4 mM (*P* = 0.0103) and CI-994 at 2.5 µM (*P* = 0.0416) and 10 µM (*P* = 0.0205) for 72 hours showed a significant increase in SSTR2 expression. **C,** QGP-1 cells treated with VPA at 4 mM for 72 hours (*P* = 0.0366) had a significant increase in SSTR2 expression. **D,** Cells treated with CI-994 had no significant increase in SSTR2 expression. **E,** QGP-1 cells treated with CI-994 at 2.5 µM (*P* = 0.0159) and 5 µM (*P* = 0.0174) for 5 days showed a significant increase in SSTR2 expression. **F,** GOT-1 cells treated with CI-994 at 5 µM (*P* = 0.0051) and 10 µM (*P* < 0.0001) for 72 hours had a significant increase in SSTR2 expression. Data are expressed as mean ± SEM, (*n* = 3). \**P* < 0.05, \*\**P* < 0.01, \*\*\**P* < 0.001, \*\*\*\**P* < 0.0001.

### HDACi treatment increases SSTR2 surface protein expression in a xenograft model expressing low basal SSTR2 levels

Using a flank xenograft model, we initially performed pilot studies in both BON-1 and QGP-1 tumors using CI-994 treatment for 3, 6, or 10 days, to determine the best SSTR2 surface response, depending on duration of drug exposure. We found a trend towards higher SSTR2 surface expression in QGP-1 tumors treated with CI-994 treatment for 10 days at both 5 and 10 mg/kg (Fig. 4A and B). We demonstrated low basal SSTR2 expression in QGP-1 control tumors compared to BON-1 control tumors (Supplementary Fig. 2A and B), and therefore chose to continue our subsequent experiments using the QGP-1 xenograft model as a representative model of human low- to negative-SSTR2–expressing NETs. Similar to our FACS results, we showed increased SSTR2 expression by IF staining after CI-994 treatment in QGP-1 xenograft tumors compared to the controls (Supplementary Fig. 2B and C). Immunohistochemistry (IHC) staining for chromogranin A was performed to confirm a NET origin in QGP-1 tumors (Supplementary Fig. 2D). A Ki67 proliferation index was also perfomed (Supplementary Fig. 2E), which revealed a high Ki67 index. These results are consistent with QGP-1 representing a model for high-grade, low-SSTR2–expressing tumors.

**Figure 4.**
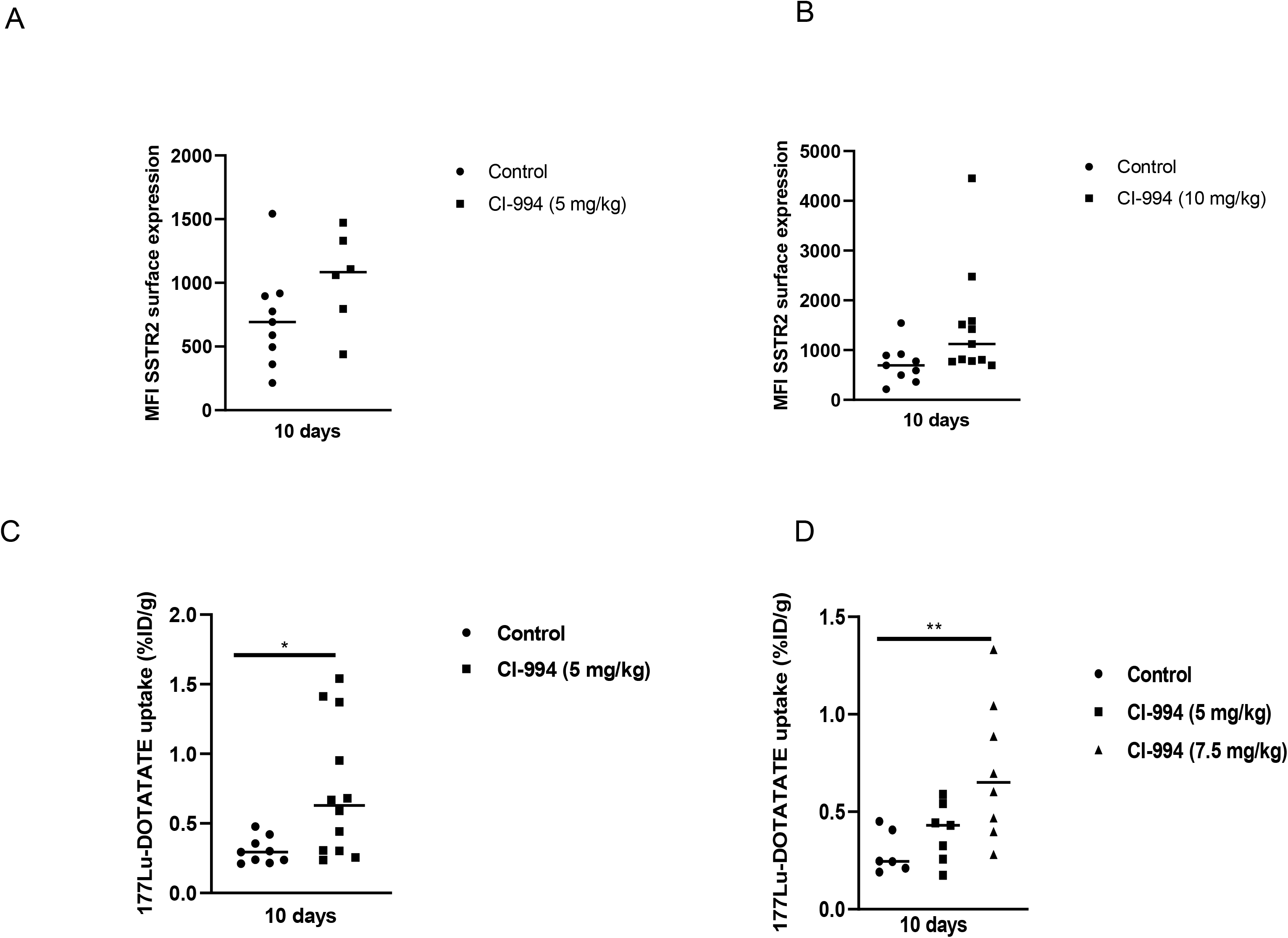
HDACis increase SSTR2 surface expression and ^177^Lu-DOTATATE uptake *in vivo.* **A,** There was an increase in SSTR2 expression with CI-994 (5 mg/kg) after 10 days of treatment, though the change was not significant (control *n* = 9, treated *n* = 6). **B,** There was an increase in SSTR2 expression with CI-994 (10 mg/kg) after 10 days of treatment, though the change was not significant (control *n* = 9, treated *n* = 11). **C,** Higher ^177^Lu-DOTATATE uptake in QGP-1 tumors indicates increased SSTR2 expression and increased therapeutic potential. A significant increase of ^177^Lu-DOTATATE was observed in tumors of mice treated with CI-994 for 10 days at 5 mg/kg (*P* = 0.0175, control *n* = 9, treated *n* = 12). **D,** Persistent uptake of ^177^Lu-DOTATATE after a 72-hour interval without CI-994 was found, with a significant increase observed in mice treated with CI-994 at 7.5 mg/kg for 10 days (*P* = 0.0092, control *n* = 6, CI-994 (5 mg/kg) *n* = 7, CI-994 (7.5 mg/kg) *n* = 8. Each circle, square, or triangle represents one tumor. Data are expressed as mean ± SEM, \**P* < 0.05, \*\**P* < 0.01.

Importantly, we did not find any significant changes in SSTR2 surface upregulation with another HDACi, VPA, at any timepoint in our *in vivo* studies (3, 6, or 10 days of treatment). Therefore, we did not use this drug for further clinical investigation (Supplementary Fig. 2F, shown at 10 days of treatment). Body weight changes shown in Supplementary Fig. 2G and H correspond to *in vivo* studies shown in Fig. 4A and B. These demonstrated minimal toxicity of CI-994 treatment in mice undergoing 10 days of daily i.p. treatment at 5 mg and 10 mg/kg, when compared with the control group.

### CI-994 treatment increases ^177^Lu-DOTATATE uptake within QGP-1 xenograft tumors

To assess whether *in vitro* increases in both SSTR2 expression and plasma membrane quantity translate to increased uptake *in vivo*, we treated QGP-1–engrafted mice with CI-994 and quantified ^177^Lu-DOTATATE uptake in tumors. We found a significant increase in ^177^Lu-DOTATATE uptake in tumors treated with CI-994 for 10 days (Fig. 4C). Our findings confirm increased cell-surface SSTR2 after HDAC inhibition that resulted in increased accumulation of ^177^Lu-DOTATATE in QGP-1 tumors, and confirmed the potential of this approach. Similarly, we found a significant increase in ^177^Lu-DOTATATE uptake in tumors pretreated with CI-994 (7.5mg/kg) after a-72-hour interval without additional CI-994 treatment. This indicates the potential to persistently upregulate SSTR2 expression for longer periods of time, even after removal of the drug, and thus continuously improve tumor uptake of ^177^Lu-DOTATATE (Fig.4D). However, this ^177^Lu-DOTATATE uptake was not found to be significant at the lower dose of CI-994 (5 mg/kg). The tumor-to-organ ratio of ^177^Lu-DOTATATE uptake showed at least a 2-fold increase in CI-994 (5 mg/kg)-treated QGP-1 tumors compared to control tumors, which represents a key feature for imaging contrast in SSTR2-targeted PRRT (Supplementary Fig. 3A and D). Body weight changes in the corresponding *in vivo* studies (Fig. 4C and D), which demonstrated limited HDACi toxicity, are shown in Supplementary Fig. 3B and E. The individual changes in tumor volume in the above studies are shown in Supplementary Fig. 3C and F, showing identical tumor sizes at the start of HDACi treatment, with no significant differences between treatment and control groups. This indicates a minimal effect of HDACi alone on tumor growth within the 10 days of i.p. injections.

### CI-994 pretreatment combined with ^177^Lu-DOTATATE promotes tumor regression in an SSTR2-deficient xenograft model compared to standard therapy

Using an identical 10-day CI-994 pretreatment model, the mice that received a single intravenous administration of 30 MBq of ^177^Lu-DOTATATE after CI-994 pre-treatment demonstrated a significant reduction in tumor growth compared to the control group (*P* < 0.0001) and to the group receiving standard therapy, i.e. 30 MBq ^177^Lu-DOTATATE alone (*P* = 0.0028) (Fig. 5A). This was confirmed 11 and 15 days after ^177^Lu-DOTATATE injections. And, the effects of combination therapy were additive, i.e. there was no interaction effect between CI-994 and ^177^Lu-DOTATATE. The clear difference in tumor growth after 5 days of ^177^Lu-DOTATATE therapy between the two CI-994–pretreated groups signaled a strong response to ^177^Lu-DOTATATE. Tumor growth was not slowed in the ^177^Lu-DOTATATE–only treatment group, potentially due to SSTR2 deficiency. Notably, the combined CI-994 and ^177^Lu-DOTATATE treatment appeared well-tolerated, with limited toxicity as evidenced by minimal changes in mouse body weight (Fig. 5B). Tumors of mice treated with combined CI-994 and ^177^Lu-DOTATATE were significantly smaller compared to tumors of mice treated ^177^Lu-DOTATATE alone (Fig. 5C). Further, Pan H3 staining revealed increased open chromatin (red foci, Pan H3) in tumors treated with CI-994 in comparison to control tumors. And increased DNA damage (green foci, γH2AX) was observed in these CI-994–pretreated tumors in comparison to control after ^177^Lu-DOTATATE therapy (Supplementary Fig. 4 A–D).

**Figure 5.**
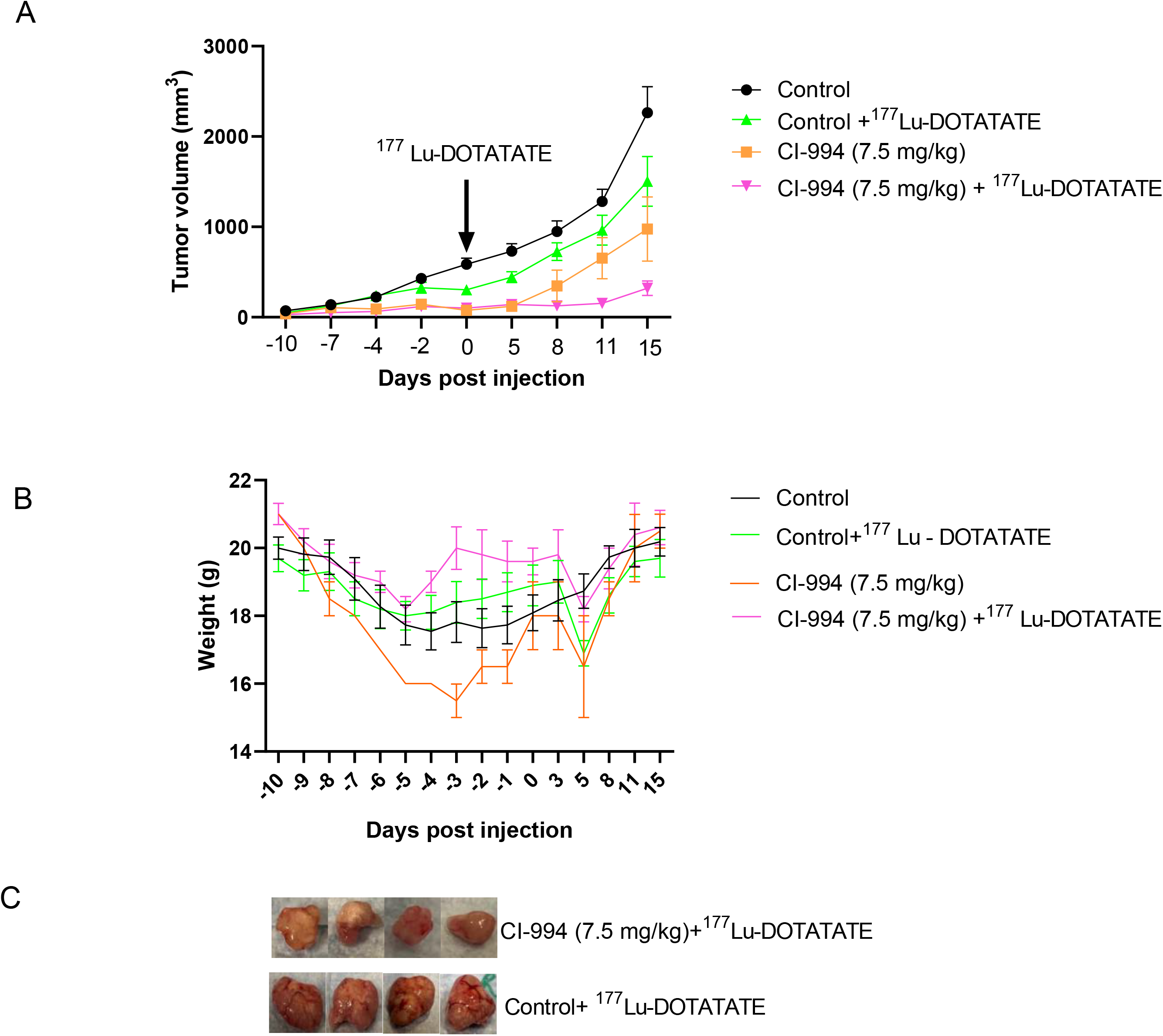
Increased ^177^Lu-DOTATATE uptake induced by CI-994 treatment promotes tumor regression in QGP-1 xenograft model. **A,** Tumor volume (mm^3^) ± SEM from start to the end of pretreatment with CI-994 and injection of 30 MBq ^177^Lu-DOTATATE at day 0 (arrow). Mice were then observed for tumor growth responses to ^177^Lu-DOTATATE for 15 days. Day 11: control vs CI-994 + ^177^Lu-DOTATATE (*P* = <0.0001); control + ^177^Lu-DOTATATE vs CI-994 + ^177^Lu-DOTATATE (*P* = 0.0008). Day 15: control vs CI-994 + ^177^Lu-DOTATATE (*P* <0.0001); control +^177^Lu-DOTATATE vs CI-994 + ^177^Lu-DOTATATE (*P* = 0.0028). **B,** Weight (g) ± SEM of mice from start to the end of treatment with CI-994 and ^177^Lu-DOTATATE. Control *n* = 18, Control +177Lu-DOTATATE n=18, CI-994 n=4, CI-994 + 177Lu-DOTATATE n=10. Data are expressed as mean ± SEM. **C,** Representative images of QGP-1 xenograft tumors collected after euthanasia, comparing the combination CI-994 +^177^Lu-DOTATATE therapy group to the control + ^177^Lu-DOTATATE group.

## Discussion

Patients with high-grade, SSTR2-negative PNETs are refractory to PRRT, with poor responses to chemotherapy (7) and are without other therapeutic options. Thus they represent an unmet challenge. Because patients with low-grade, SSTR2-positive metastatic NETs have shown significant progression-free survival benefits in response to SSTR2-targeted PRRT (9), we propose to improve tumor responsiveness to PRRT in patients with high-grade, receptor-negative metastatic PNETs by increasing their SSTR2 levels.

In our study, we demonstrate for the first time the clinical relevance of using ^177^Lu-DOTATATE as a targeted therapy in a receptor-deficient *in vivo* PNET model. Our ultimate goal was to discover and characterize novel, SSTR2-directed, epidrug-based treatments to improve responses to PRRT. Changes in epigenetic marks have been demonstrated in various cancers, including PNETs (33, 34), and because these changes are reversible, they serve as targets for the modulation of gene expression. We found that higher NET grades correlated with increased SSTR2 promoter methylation levels in an NIH NET patient cohort, which provides an explanation for the silenced expression of SSTR2, as shown in other datasets (35). In high-grade PNETs, SSTR2 seems lost due to epigenetic gene-silencing mechanisms (14,36–38); and a few studies have reported a reversal of these mechanisms by treating NET cell lines with DNMTis and HDACis (39), thus inducing DNA hypomethylation and histone acetylation, respectively (34,40,41).

In our *in vitro* studies, we observed that QGP-1, BON-1, and GOT-1 cell lines have different baseline expression levels of SSTR2 (QGP-1 < BON-1 < GOT-1), consistent with prior reports (26). To the best of our knowledge, we are the first to show that higher DNA methylation correlates with lower SSTR2 expression in these three NET cell lines. We demonstrated this by using epidrug treatments to increase SSTR2 expression, eliciting a particularly strong response in QGP-1, which harbors hypermethylated SSTR2 promoter levels and low SSTR2 expression. Accordingly, we judged QGP-1 to be most representative of the high-grade, SSTR2-negative human PNETs and used it for our *in vivo* therapeutic studies. Examining SSTR2 cell surface expression, we demonstrated a significant and strong dose-dependent upregulation at 72 hours and at 5 days of HDACi treatment, but not at shorter timepoints. These results indicate that both time course and dose play crucial roles in regulating SSTR2 transcription using epidrug-based treatments. Others have reported similar results for upregulation of SSTR2 mRNA in BON-1 and GOT-1 with HDACi treatments (12,14,38). In an *in vitro* study in BON-1 and NCI-H727 cells (14), the reversibility of *SSTR2* gene expression upregulation after HDACi (VPA and CI-994) withdrawal returned to baseline within 24 hours. *In vivo*, our results confirm that effects induced by low doses of HDACi treatment (5 mg/kg) for a short duration were largely and rapidly reversible. Interestingly, the higher dose of CI-994 (7.5 mg/kg) led to persistent upregulation of SSTR2 expression, even after withdrawal of the drug for 3 days, showing a significant increase in ^177^Lu-DOTATATE uptake within tumors.

To the best of our knowledge, our study is the first to report *in vivo* therapeutic results with ^177^Lu-DOTATATE in a PNET model harboring low basal SSTR2 expression. Our results demonstrate significant upregulation of SSTR2 surface expression and ^177^Lu-DOTATATE uptake after HDACi treatment in QGP-1 tumor xenografts. Most relevant was the significant tumor regression seen in HDACi-pretreated mice in response to the ^177^Lu-DOTATATE therapy, when compared to both control mice and ^177^Lu-DOTATATE-only treated mice. Importantly, this combined, sequential therapeutic effect of HDACi CI-994 and ^177^Lu-DOTATATE therapy was additive. Furthermore, the ^177^Lu-DOTATATE-only treatment group had a poorer response than the CI-994-only treatment group. We hypothesize that this may be due to the low basal SSTR2 expression in our QGP-1 xenograft model, which would explain the poor response to treatment with ^177^Lu-DOTATATE alone. This *in vivo* model is thus representative of the poor response to PRRT in humans with high-grade, receptor-negative NETs. We observed extensive DNA damage with the uptake of ^177^Lu-DOTATATE into tumors, with a higher absorbed dose translating into increased DNA damage and leading to better outcomes (combined treatment group); as shown by using γH2AX as a marker for the biological effect of ^177^Lu-DOTATATE PRRT (42).

The major strength of our study is its translational relevance, as the therapeutic efficacy of PRRT has only been proven for models characterized by high baseline SSTR2 surface expression levels. Our results suggest that receptor-deficient tumors, such as in our QGP-1 model, may respond to PRRT using ^177^Lu-DOTATATE when pretreated with selective epidrugs. These epidrugs will increase SSTR2 expression, thus expanding the bounds of PRRT clinical efficacy for use in treating patients with high-grade SSTR2-negative PNETs.

In line with our *in vivo* studies, these epidrug treatments have been demonstrated to be safe in patients, as shown in a clinical imaging study analyzing the safety of the HDACi vorinostat (43). However, more studies are necessary to understand the toxicity profiles in patients when these epidrugs are combined with ^177^Lu-DOTATATE therapy.

Our study has a few limitations. Unfortunately, there are no representative high-grade metastatic NET models for GEP-NETs. We utilized three NET cell lines for our *in vitro* studies to best represent and cover all possible responses to our investigational drugs due to varying degrees of SSTR2 surface expression. To overcome this limitation in our *in vivo* studies, we selected the QGP-1 model, shown to harbor the lowest SSTR2 surface expression, so as to best represent human high-grade, receptor-negative NET behavior.

In conclusion, our preclinical data demonstrate that pretreatment with the HDACi CI-994 improves ^177^Lu-DOTATATE therapy compared to PRRT alone for the treatment of SSTR2-deficient tumors. Using epidrug-based treatment regimens to elicit an increase in SSTR2 surface expression in patients with metastatic PNETs, we intend to improve their tumor responsiveness to PRRT. Our study thus forms the basis for a clinical trial testing the therapeutic efficacy of HDACi CI-994 pretreatment in combination with ^177^Lu-DOTATATE therapy in patients with high-grade, SSTR2-negative metastatic PNETs.

## Declarations

## Acknowledgements

This work was supported by NCI/NIH Intramural Funding to Samira Sadowski ZIA BC 011899. FEE was funded by ZIA BC 011800 and ZIA BC 010891.

Special thanks to Dr. Peter Choyke, Chief of the Molecular Imaging Branch, for his support and the use of his radiation/PET imaging facilities.

We thank the flow cytometry core facility, National Heart, Lung, and Blood Institute (NHLBI), NIH for their help with flow cytometry experiments, and the CCR Microscopy Core Facility, National Cancer Institute (NCI) for their help with microscopy imaging.

We thank Dr. Sunita Agarwal (NIDDK) for her critical review of the paper.

We thank Dr. Paden King, Dr. Noriko Sato, Neil Alilin Aian, and Collen Olkowski for their technical support. We thank Dr. Joanna Shih, Biostatistics Branch, for her statistical support.

## Authors’ contributions

SS designed the study. RS, BE, JM, and KB conducted the experiments. RS, FE, and SS analyzed the data. RS, MZ, JM, and SS wrote the manuscript. All authors read and approved the final manuscript.

## Ethics approval

All procedures were carried out in accordance with the NIH IC Animal Care & Use Committee guidelines, as well as Institutional radiation guidelines, and were approved by the NIH Radiation Safety Committee, Bethesda, Maryland USA.

## Consent for publication

Not applicable

## Availability of data and material

The datasets generated and/or analyzed during the current study are available from the corresponding author upon reasonable request.

