## Supplemental Material for "Upregulation of Somatostatin Receptor Type 2 in a Receptor-Deficient *In Vivo* Pancreatic Neuroendocrine Tumor Model Improves Tumor Response to Targeted ^177^Lu-DOTATATE"

### Supplementary Figure Legends

**Supplementary Figure 1.** Validation of a commercially available SSTR2 antibody. **A,** qRT-PCR was performed on separate BON-1 PNET cell clones stably expressing either a non-targeting (NT) shRNA or pre-validated human SSTR2-specific shRNA (#64-shRNA or #87-shRNA). Stable knockdown of SSTR2 mRNA was demonstrated for both SSTR2-specific shRNA: ~75% knockdown seen in #64-shRNA cells and ~65% knockdown seen in #87-shRNA cells. **B,** Protein lysates from the three separate shRNA-expressing BON-1 cell clones were resolved by SDS-PAGE and probed for SSTR2 using a commercially available SSTR2-specific antibody. Lysates were further probed for β-actin, as a loading control, to confirm equal loading of protein. Using an SSTR2-specific antibody from Boster Bio (# M01689), decreased expression of a specific band at roughly 75 kDa was noted in cells expressing both #64-shRNA and #87-shRNA, with relative knockdown trends similar to results determined by qRT-PCR.

**Supplementary Figure 2.** *In vivo* xenograft protein expression. **A,** SSTR2 surface expression (red) in BON-1 control tumors, $n = 4$. **B,** SSTR2 surface expression (red) in QGP-1 control tumors, $n = 4$. **C,** Surface expression of SSTR2 (red) with CI-994 treatment. DAPI was used to stain cell nuclei (blue), $n = 4$ **D,** Chromogranin A staining (brown) to confirm NET origin. **E,** Ki67 staining (brown) to confirm proliferation index (scale bar = 100 μm). **F,** No change in SSTR2 surface expression in mice treated with VPA for 10 days (control $n = 4$; VPA 300 mg/kg, $n = 8$). **G,** Weight of mice from start to the end of treatment with CI-994 (5 mg/kg) for 10 days. **H,** Weight of mice from start to the end of treatment with CI-994 (10 mg/kg) and control group, corresponding to studies illustrated in **Fig. 4A** and **B**.

**Supplementary Figure 3.** *In vivo* biodistribution studies and mouse weights. **A**, QGP-1 tumors (CI-994, 5 mg/kg): biodistribution in blood and organs (heart, lungs,liver, spleen, kidney, stomach, small and large intestine, muscle, femur) in control and CI-994–treated tumors, collected 24 hours after $^{177}$Lu-DOTATATE injection. Measured as the tumor-to-organ ratio of $^{177}$Lu-DOTATATE uptake in CI-994–treated QGP-1 mice compared to control mice. **B,** Weight of mice from start to the end of CI-994 treatment from *in vivo* $^{177}$Lu-DOTATATE uptake study illustrated in **Fig. 4C**. **C,** Tumor volumes from start to the end of CI-994 treatment from same study. **D,** QGP-1 tumors (CI-994, 5mg/kg and 7.5 mg/kg): biodistribution in blood and organs (heart, lungs,liver, spleen, kidney,stomach, small and large intestine, muscle, femur) in control and CI-994–treated tumors, collected 24 hours after $^{177}$Lu-DOTATATE injection. Measured as the tumor-to-organ ratio of $^{177}$Lu-DOTATATE uptake in CI-994 treated QGP-1 mice compared to control mice. **E,** Weight of mice from start to the end of CI-994 treatment from *in vivo* $^{177}$Lu-DOTATATE uptake study illustrated in **Fig. 4D**. **F,** Tumor volumes from start to the end of CI-994 treatment from the same study.

**Supplementary Figure 4.** Correlative studies of protein expression in QGP-1 xenografts. **A,** IF-stained slides for the expression of Pan-acetylated H3 (red) in control mice ($n$ = 4). **B,** Expression of Pan-acetylated H3 in mice treated with CI-994 (7.5 mg/kg) ($n$ = 4). **C,** IF-stained slides for the expression of γH2AX (green) in $^{177}$Lu-DOTATATE-only treated mice ($n$ = 4). **D,** Expression of γH2AX in mice pretreated with CI-994 (7.5 mg/kg) after 15 days of treatment with $^{177}$Lu-DOTATATE ($n$ = 4). DAPI was used to stain cell nuclei (blue) (20X, scale bar = 50 µm, 63X and 100X scale bar = 5 µm).

**Supplementary Table 1: Antibodies used in this study.**

A

B

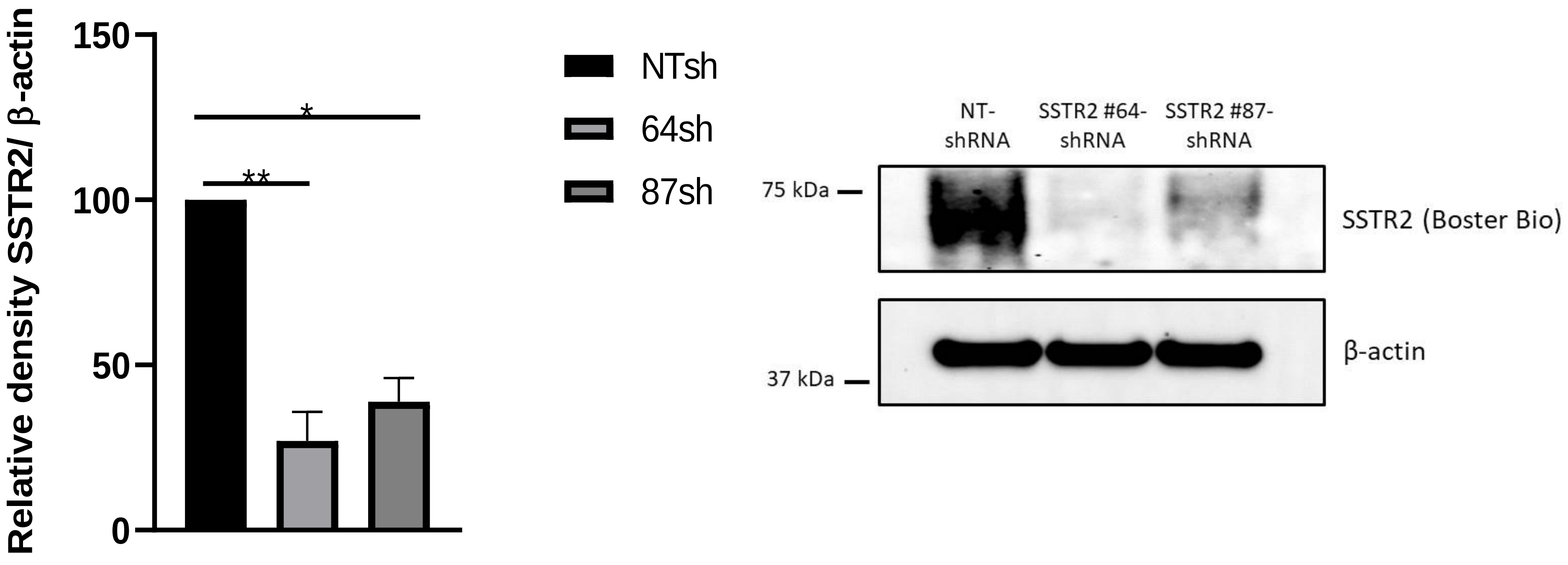


Supplementary Figure 1

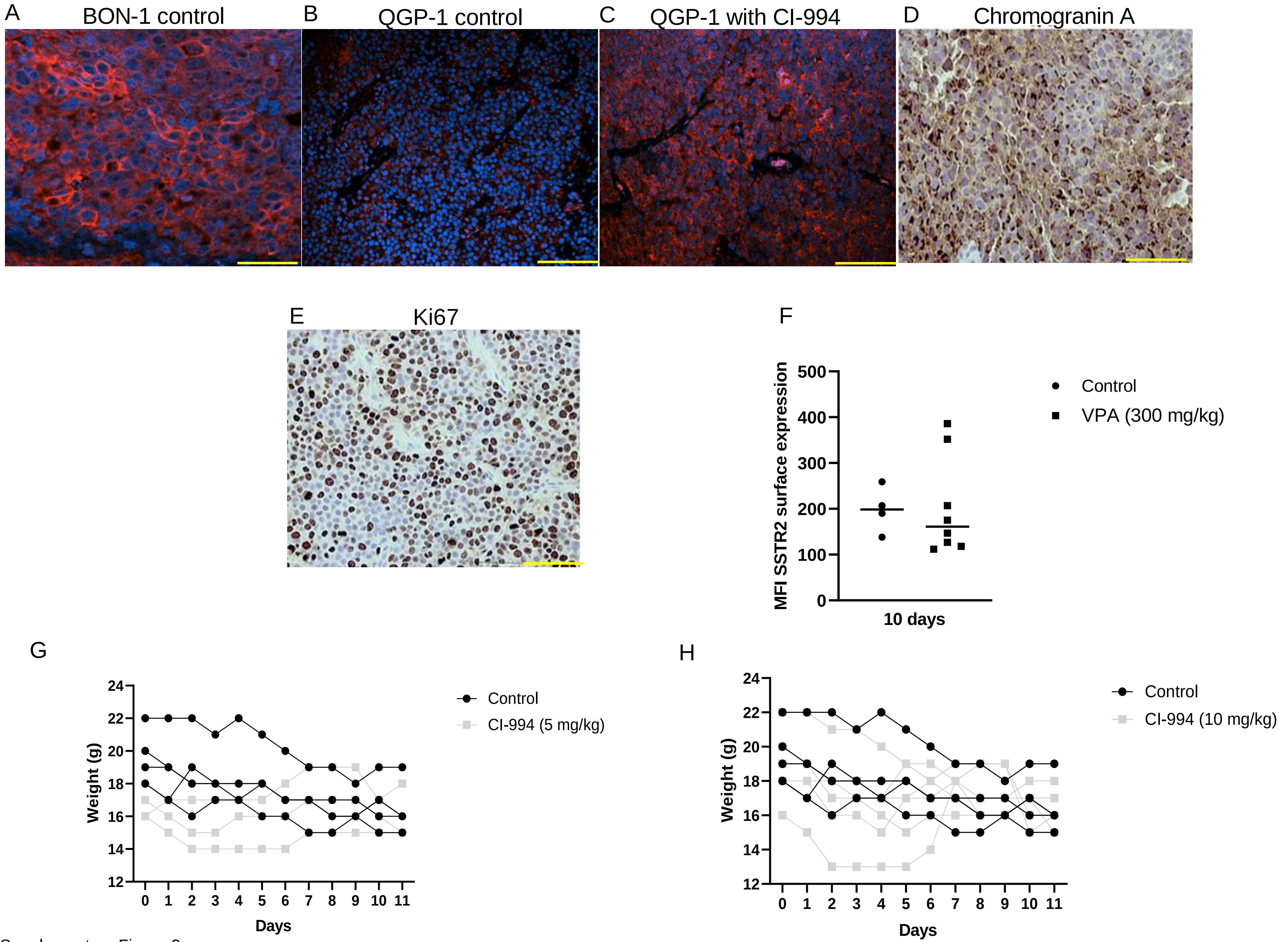


Supplementary Figure 2

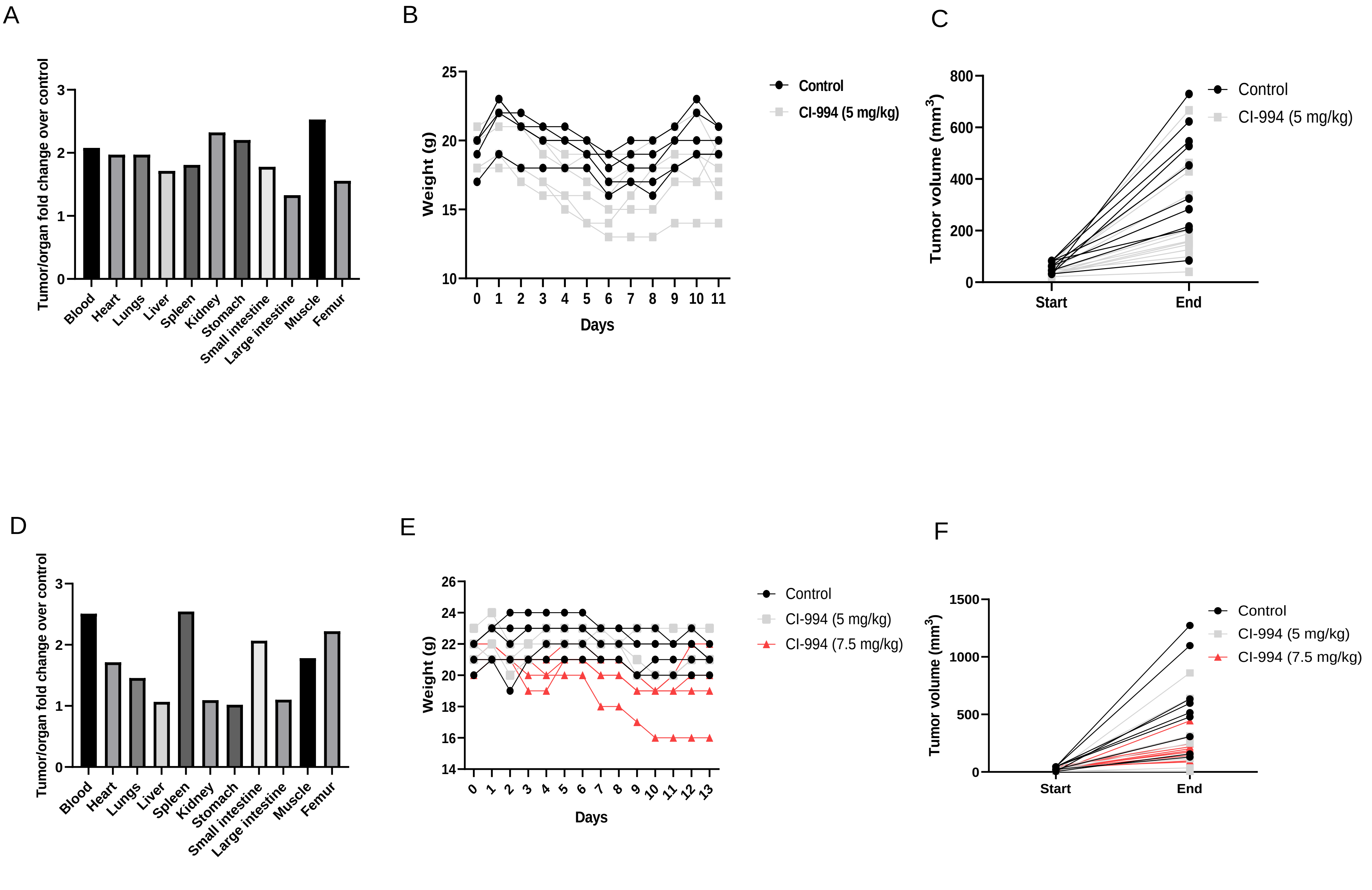


Supplementary Figure 3

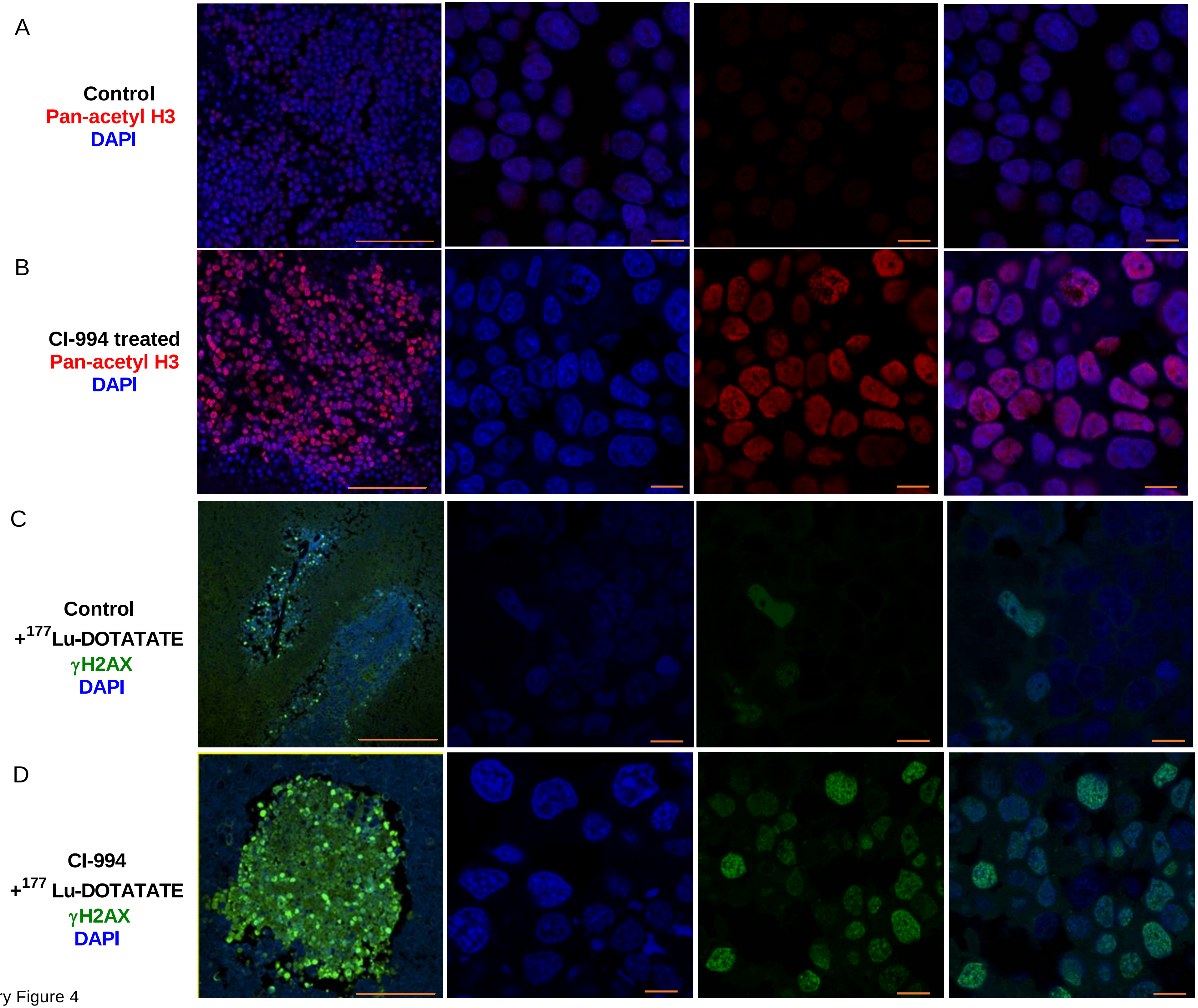


Supplementary Figure 4

**Table 1**

| Antibody | Catalog no. | Company | Dilution |
|---|---|---|---|
| SSTR2 (Western)<br><br>SSTR2 (FACS) | M01689<br><br>IC4224G | Boster Bio<br><br>R&D Systems | 1:1000<br><br>1µg/5ul |
| GAPDH | 2118 | Cell Signaling | 1:2000 |
| Histone 3 (H3) | 9715L | Cell Signaling | 1:500 |
| Pan-acetylated H3 | MAB397 | Millipore | 1:500 |
| HRP-tagged anti mouse | 7076P2 | Cell Signaling | 1:5000 |
| HRP-tagged anti rabbit | 7074P2 | Cell Signaling | 1:5000 |
| Chromogranin (IHC) | NB-120-15160. | Novus | 1:50 |
| Ki67 (IHC) | ab16667 | Abcam | 1:200 |
| SSTR2 (IF) | UMB1 - ab134152- | Abcam | 1:25 |
| Pan-acetylated H3 (IF) | 06-599 | Millipore | 1:50 |
| γH2AX (IF) | 05-636 | Millipore | 1:800 |

Supplementary Table 1
